# Coupled Solvent and Protein Dynamics Confer Differences in Exon-19 Deletion Mutants of the Epidermal Growth Factor Receptor Kinase

**DOI:** 10.1101/2024.10.04.616731

**Authors:** Keshav Patil, Debdas Dhabal, Kumar D. Ashtekar, Yuko Tsutsui, Krishna Suresh, Hrishabh Singh, Mark A. Lemmon, Ravi Radhakrishnan

## Abstract

Deletions in Exon-19 of the epidermal growth factor receptor (EGFR) play a pivotal role in the pathogenesis of non-small cell lung cancer (NSCLC), influencing patient response to tyrosine kinase inhibitors (TKIs). Although these mutations are known to affect treatment efficacy, the precise molecular mechanisms have been unclear. Building upon recent insights from the study [DOI: 10.1038/s41467-022-34398-z], which identified two distinct mutation profiles associated with differential drug sensitivity and clinical outcomes, our research delves into the molecular dynamics that drive these variances.

We employed molecular dynamics simulations, enhanced sampling methods, and machine learning to classify Exon-19 deletion mutations into two profiles based on their conformational dynamics. Profile 1 mutations display only localized motions in key subdomains in their fluctuations about the equilibrium state, and a high affinity for ATP and consequent resistance to TKIs, while profile 2 mutations show reduced ATP binding affinity due to delocalized motion characterized by an increased flexibility between the N- and C-lobes of the EGFR kinase domain. This structural flexibility perturbs the ATP binding site, leading to decreased affinity and, heightened sensitivity to TKIs.

Our use of the INDirect Umbrella Sampling (INDUS) technique has shed light on the collective solvent dynamics, further elucidating the coupling between long timescale solvent fluctuations and protein conformational dynamics, that likely contributes to the observations in HDX-MS studies. Our free energy analysis, covering timescales relevant to both HDX-MS and ligand interaction, provides a deeper understanding of the relationship between protein and solvent dynamics and their collective impact on drug efficacy in NSCLC with EGFR Exon-19 deletions.

**Significance Statement:** EGFR Exon 19 deletion mutations are key drivers in non-small cell lung cancer (NSCLC), yet their drug sensitivity to tyrosine kinase inhibitors (TKIs) varies significantly. This study identifies two mutation profiles: mutations that exhibit high ATP binding affinity and localized conformational motion, driving TKI resistance, and mutations that show reduced ATP affinity due to delocalized structural flexibility, enhancing TKI sensitivity. Using molecular dynamics simulations and free energy sampling techniques, we reveal how solvent fluctuations and protein dynamics collectively affect drug binding and efficacy. These findings provide a mechanistic basis for differential drug sensitivity, informing precision medicine strategies for NSCLC patients with EGFR Exon-19 mutations.

## Introduction

Tyrosine kinase inhibitors (TKIs) are the foremost treatment option for EGFR mutations (1–3) in non-small cell lung cancer (NSCLC). Exon-19 and Exon-21 mutations represent the majority, over 90%, of the oncogenic EGFR alterations (4–6). The most common Exon 19 mutation, constituting 33% of EGFR mutations in NSCLC, is a deletion of five amino acids (delE746-A750) in the loop that joins the third β-strand of the EGFR tyrosine kinase domain (TKD) with the vital αC helix. However, many other rarer Exon 19 deletions are seen in NSCLC patients and appear to be associated with reduced sensitivity to EGFR TKIs (7, 8). These EGFR Exon 19 deletion mutations differ in the length of their deletion lengths, and many are indels. Recent studies indicate that the differential drug sensitivity of Exon 19 variants to various EGFR inhibitors reflects their conformational dynamics (9). Comparison of a set of Exon 19 deleted variants suggested that their drug binding affinity (with erlotinib) was essentially unchanged, but they showed alter *K*_M_ values for ATP, suggesting that variation in ATP binding affinity is the main determinant of (ATP-competitive) drug sensitivity. Grouping the variants studied according to their ATP binding affinity and analyzing their structural dynamics through HDX-MS to acquire exchange profiles allowed two distinct groups or profiles to be identified. Profile 1 mutations (e.g., L747P, del747-750-insP) showed higher ATP binding affinity, translating to a lower *K*_M_, whereas Profile 2 mutations (e.g., del746-750, del-747-753-insS) exhibited increased flexibility and disorder in the ATP binding site, resulting in reduced ATP binding affinity. This variable ATP binding affinity contributed to drug resistance due to the heightened competition between the kinase inhibitor and ATP for the binding site. Kaplan-Meier survival analyses further indicated that patients with Profile 1 mutations had a shorter median progression-free survival on erlotinib treatment compared to those with Profile 2 mutations. We note that a similar observation between ATP binding affinity and drug sensitivity was also reported in the most common EGFR TKD mutation, L858R, in the context of an allosteric inhibitor (10).

Our aim in this paper is to use molecular dynamics (MD) simulations to delve into the molecular underpinnings of the differential dynamics in exon 19 deletion mutants. We attribute the differential TKI sensitivity of Exon 19-mutated variants to altered protein conformational dynamics, backbone hydrogen bond networks, solvent accessibility, and electrostatic interactions. We propose using MD as a cohesive framework to decode both the hydrogen deuterium exchange profiles and the structural origins of this exchange, as a critical step in comprehending how mutations affect protein function, particularly in kinases. Protein dynamics are crucial to their function and biomolecular interactions. Hydrogen-deuterium exchange mass spectrometry (HDX-MS) is an effective method for probing protein dynamics. The well-established two-step model of HDX, proposed by Linderstrom-Lang (11), is utilized to interpret the protein dynamics observed, where the protein’s amide N-H hydrogen transitions from a closed to an open state before exchanging with solvent deuterium. HDX-MS, in conjunction with mass spectrometry, is particularly advantageous for studying larger protein complexes without size restrictions, enhancing the molecular understanding of the dynamics involved in kinases and the impact of mutations on these dynamics. HDX provides insights into protein dynamics through changes in hydrogen bond networks, and some conformational changes can be ‘HDX-silent’ if they cause only minimal changes in hydrogen bonding. For instance, the rupture of Heme-Fe(III) contact in cytochrome c causes significant reorientation of a loop with little change in hydrogen bonding, and is HDX-silent (12). Correlating HDX-MS data with MD simulations has the potential to be of great value for understanding the molecular origins behind the observed deuterium exchange of biomolecules studied in HDX (13, 14).

Whereas MD simulations typically focus on the microsecond scale, HDX operates in much larger scales from seconds to hours. Accordingly, HDX can capture protein dynamics across a vast range of timescales, encompassing protein folding/unfolding, conformational transition states, ligand binding, and other biomolecular interactions. To use MD to link HDX results to their molecular origins, therefore, the timescale of MD must be expanded, which can be achieved using enhanced sampling techniques. However, implementing such techniques requires prior knowledge of the collective variables that govern protein transitions, which can be difficult to predict. To extend MD analysis in the context of HDX-MS, we apply Indirect Umbrella Sampling (INDUS), a non-Boltzmann enhanced sampling technique used to compute hydration thermodynamics and solvent density fluctuations associated with conformational transitions (15–17). This allows us to simulate the coupling between the collective solvent fluctuations and protein conformational dynamics to gain insight into long-timescale fluctuations in protein conformations in the EGFR Exon-19 deletion mutant systems. In this report, we combine MD simulations, machine learning, and HDX-MS to elucidate the impact of EGFR Exon-19 deletion mutations on short timescale (microsecond) kinase conformational dynamics. We demonstrate the critical role of hydrogen bonding networks in shaping these dynamics and, through the use of INDUS, reveal the intricate relationship between collective solvent dynamics and long timescale (i.e., milliseconds and larger) conformational fluctuations, thus improving the integration of MD simulations with HDX-MS data.

## Results

Alongside their analysis of TKI sensitivity of exon 19-deleted variants of the measured purified EGFR tyrosine kinase domain (TKD), see Fig. S1 A,B, van Alderwerelt van Rosenburgh *et al*. used HDX-MS to identify conformational dynamics as a possible origin for the differences, and proposed a basis for classifying uncommon exon 19 variants that may have predictive clinical value (9). They showed that native state dynamics of exon 19 variants differ for profile 1 and profile 2 variants as shown in Fig. S1C. Profile 1 (more TKI-resistant) variants (e.g. DelL747-A750InsP and L747P) showed generally lower degrees of exchange in the ATP-binding region of the TKD than profile 2 (more TKI-sensitive) variants (e.g. DelE746-A750 and DelL747-P753InsS). How the altered conformational dynamics as indicated by HDX translates to altered ATP binding affinity, however, is unclear. We chose variants in profile 1 and 2 that differ by only one amino acid to analyze by molecular dynamics in order to investigate the origin of this difference in conformational dynamics.

### Principal dynamics and cross correlations at the subdomain level correctly profile EGFR variants

We performed 4 μs MD simulations of all profile 1 and profile 2 variants studied experimentally by van Alderwerelt van Rosenburgh *et al*. as described in Methods. We first compared the MD trajectories for the profile 1 (resistant) Del747-750InsP and profile 2 (sensitive) Del746-750 systems. We performed principal component analysis (PCA) of the two MD trajectories and projected the configurations visited in the respective trajectories along the top two principal components (Fig. 1A). Based on these two-dimensional projections, we identified clusters according to the relative frequency of conformations visited; a stark difference in the PCA projections is evident. The difference can also be visualized by comparing the root mean squared deviations about the structures corresponding to the centroids of each of the clusters as shown in Fig. 1B, see also SI Movies (SI-M1 to M9).

**Figure 1.**
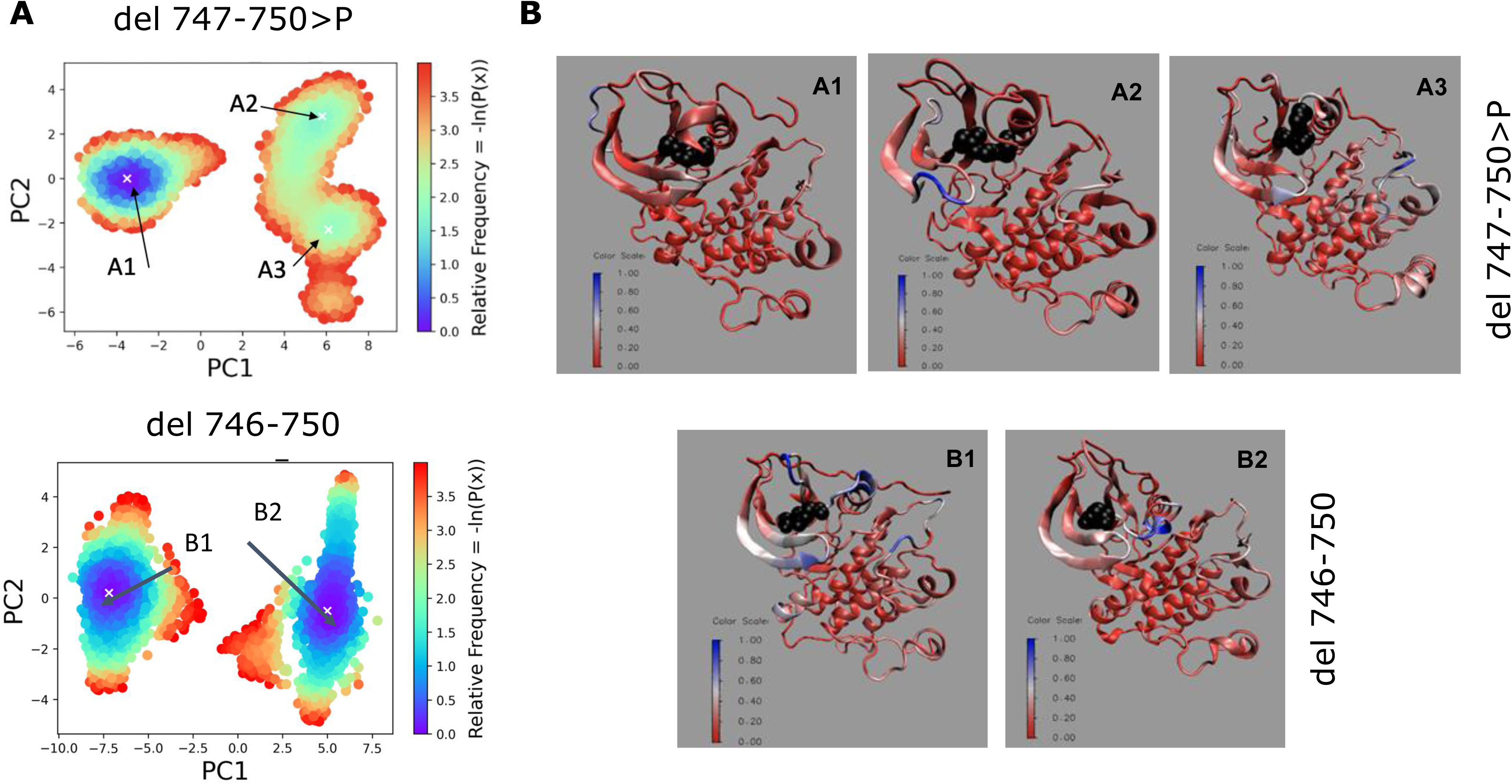
A. Projections of the conformations visited in the MD trajectories on the top two principal components with relative frequencies colored according to the color bar for a profile 1 variant: del 747-750>P and a profile 2 variant: del 747-751>P differing by just one amino acid. B. Structures extracted from the PCA clusters (A1-A3 and B1-B3) as marked in A. The color bar represents the RMSD fluctuations. The profile 1 variant predominantly shows localized motion of the activation loop and the αC helix. Whereas the profile 2 motion shows delocalized motion involving other subdomains in the C-lobe and the N-lobe.

We summarize the results of the PCA on all other variants studied experimentally in Table 1, (see also SI Movies, SI-M1 to SI-M9), wherein the principal motions in certain kinase subdomains are listed. We examine the essential motions in N-lobe and C-lobe, since their motion provides global flexibility for the ATP binding pocket. We observe that the motions in the profile 1 variants occur largely around P-loop, αC-helix and Activation loop, whereas these motions also extend to N–lobe and C-lobe regions for the profile 2 variants, see SI Movies. To provide a quantitative description and classification of these principal motions we next computed residue-wise dynamical cross correlation matrices from the atomic fluctuations, and coarse-grained these matrices to subdomain-level cross correlations as described in Methods and Table 2.

**Table 1:**
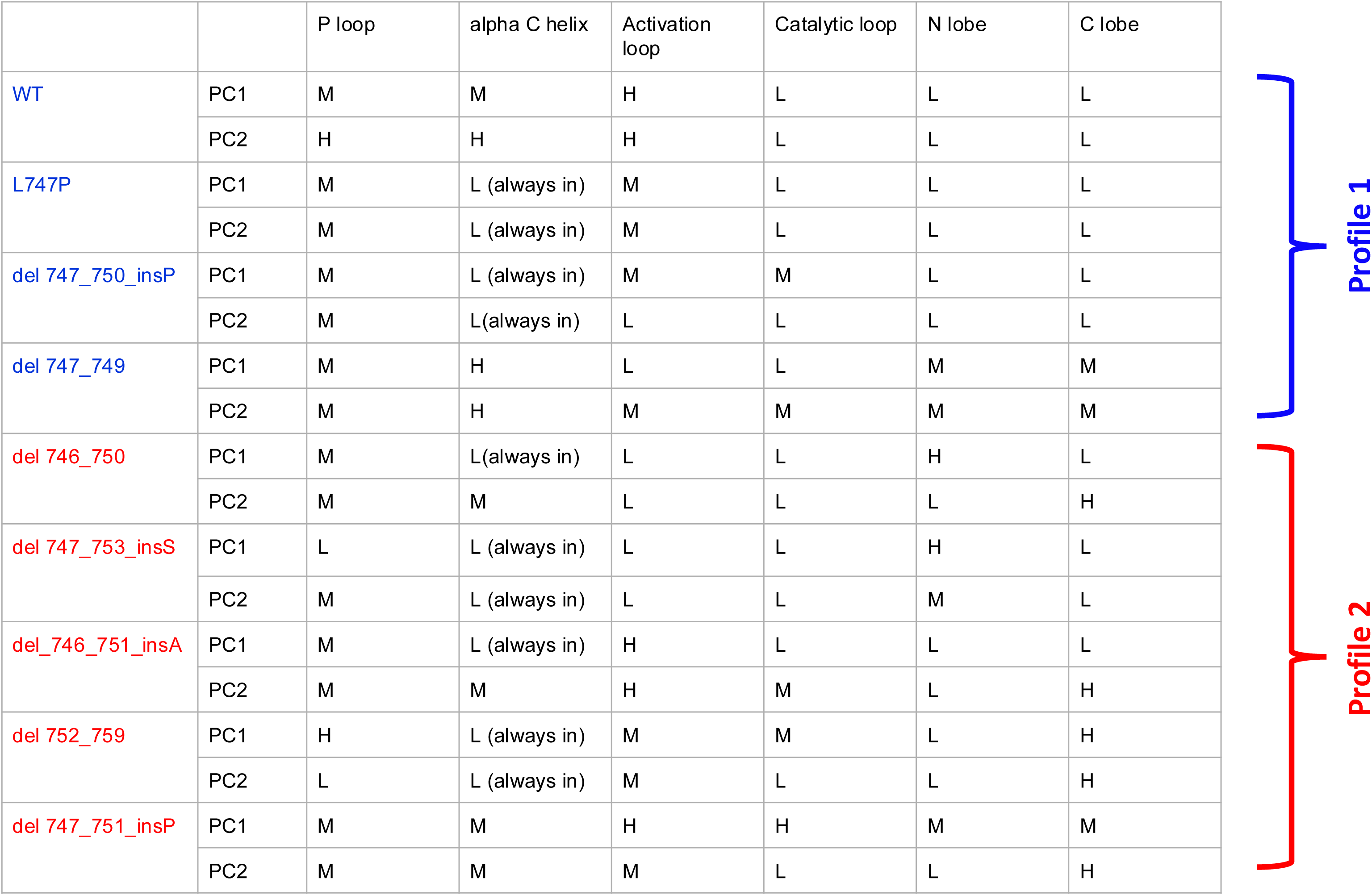
Observed motion projected along the top two principal components in several subdomains for EGFR deletion mutants classified as high (H), medium (M), and low (L).

**Table 2:**
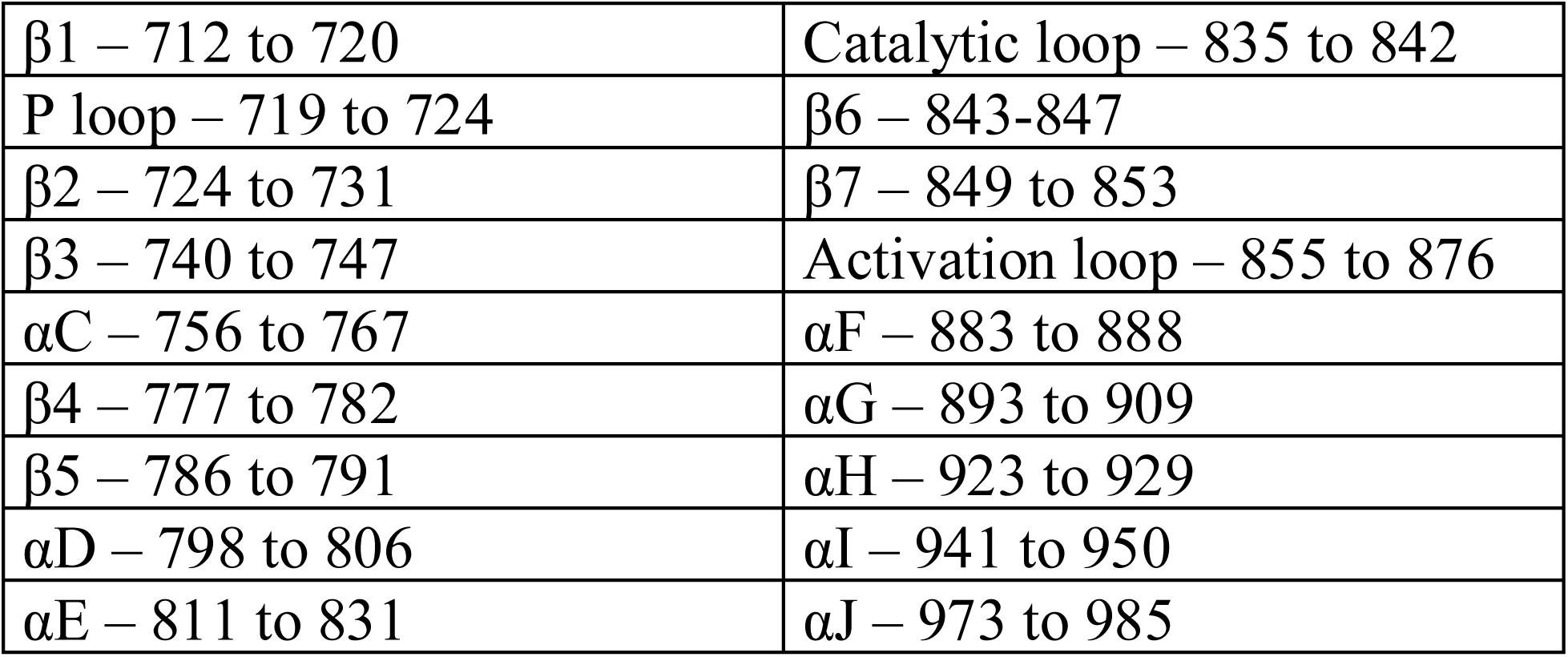
Subdomains and key sub-regions of the EGFR kinase domain. These subdomain definitions are used to compute the dynamical cross correlation matrices.

|  |  |
| --- | --- |
| $\beta 1$ – 712 to 720 | Catalytic loop – 835 to 842 |
| P loop – 719 to 724 | $\beta 6$ – 843-847 |
| $\beta 2$ – 724 to 731 | $\beta 7$ – 849 to 853 |
| $\beta 3$ – 740 to 747 | Activation loop – 855 to 876 |
| $\alpha C$ – 756 to 767 | $\alpha F$ – 883 to 888 |
| $\beta 4$ – 777 to 782 | $\alpha G$ – 893 to 909 |
| $\beta 5$ – 786 to 791 | $\alpha H$ – 923 to 929 |
| $\alpha D$ – 798 to 806 | $\alpha I$ – 941 to 950 |
| $\alpha E$ – 811 to 831 | $\alpha J$ – 973 to 985 |

The coarse-grained correlation matrices for profile 1 variants (and wild-type) are shown in Fig. 2A, and those for profile 2 variants are shown in Fig. 2B. The qualitative categories of low, medium, and high degrees of cross correlations summarized in Table 1 are quantitatively borne out in Fig. 2. The most immediately obvious difference between profile 1 and profile 2 variants in these matrices is the greater delocalization of cross-correlations in profile 2 mutants, engaging regions in both the N- and C-lobes of the TKD. Both profiles show high degrees of correlated motion primarily in the N-lobe, and cross-correlated motions between subdomains of the N-lobe (strands β1-5, and the P-loop). Profile 1 variants show little cross-correlation outside these regions, whereas profile 2 variants, notably Del747-750InsP and Del746-751InsA, show significant cross correlation involving C-lobe β-strands and α-helices as well as the Activation loop. To further categorize these differences between profile 1 and profile 2 variants, we compared the dynamical cross correlations (DCCs) by performing a pairwise distance calculation and constructed a hierarchical cluster (Fig. 2C); we find that this unsupervised clustering is remarkably efficient in classifying the EGFR variant systems into the two separate profiles with excellent agreement with what was observed in the experiments (9). The approach readily identifies the wild-type TKD as well as Del747-749, Del747-750InsP, and L747P as profile 1. Their experimental *K*_M_ values for ATP range from 12 to 23 μM (9). The Del746-750, Del752-759 and Del746-751InsP cluster differently and have significantly higher values for *K*_M, ATP_ (91 to 164 <u>μM</u>). We note that two systems Del747-753InsS and Del747-751InsP are misclassified in this analysis. We surmise that a possible reason for this misclassification is the limited sampling of conformations in molecular dynamics (spanning just 4 μs) in comparison to the much longer timescales (seconds-to-hours) over which exchange occurs in HDX experiments. We address this limitation through enhanced sampling discussed later.

**Figure 2.**
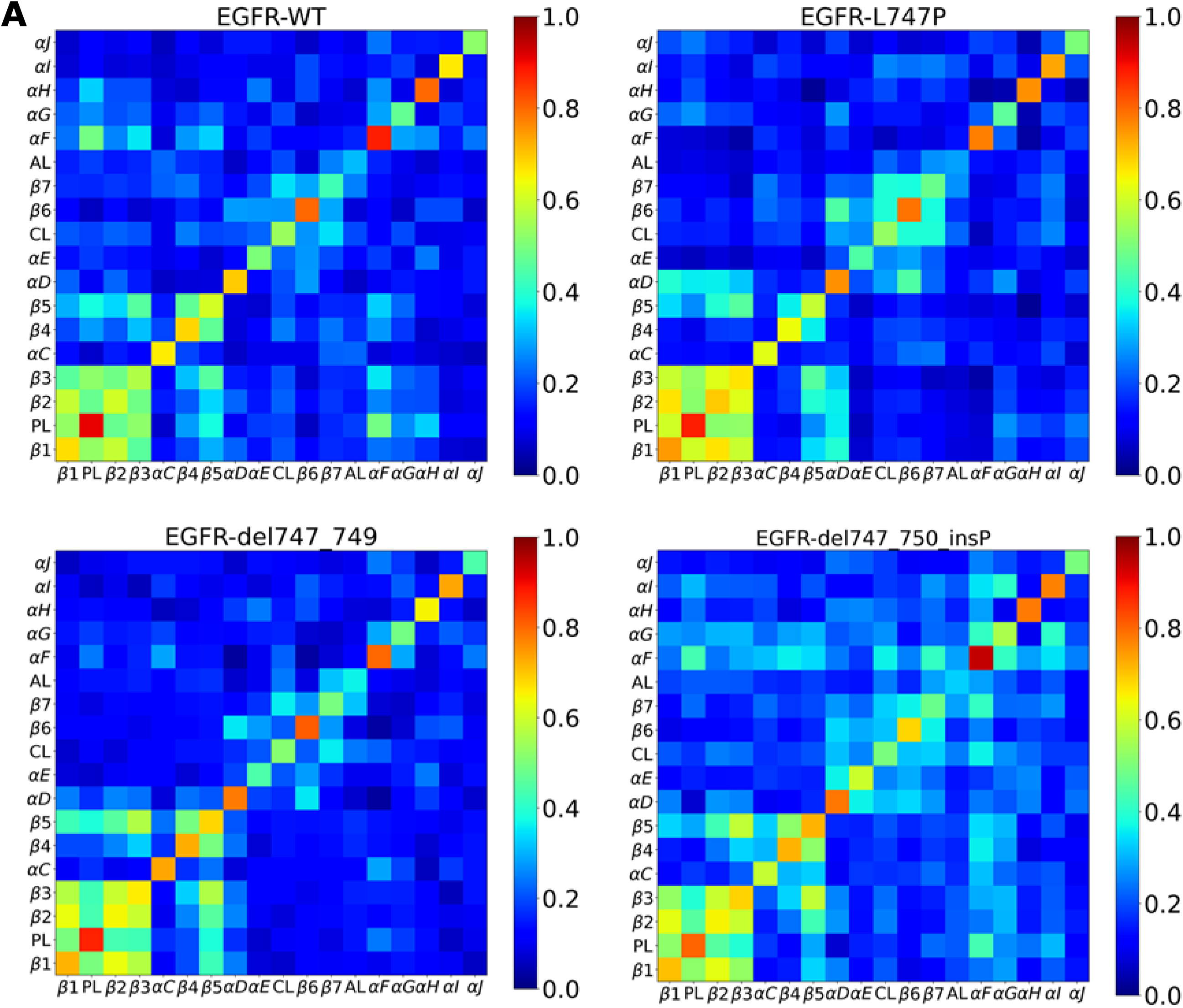
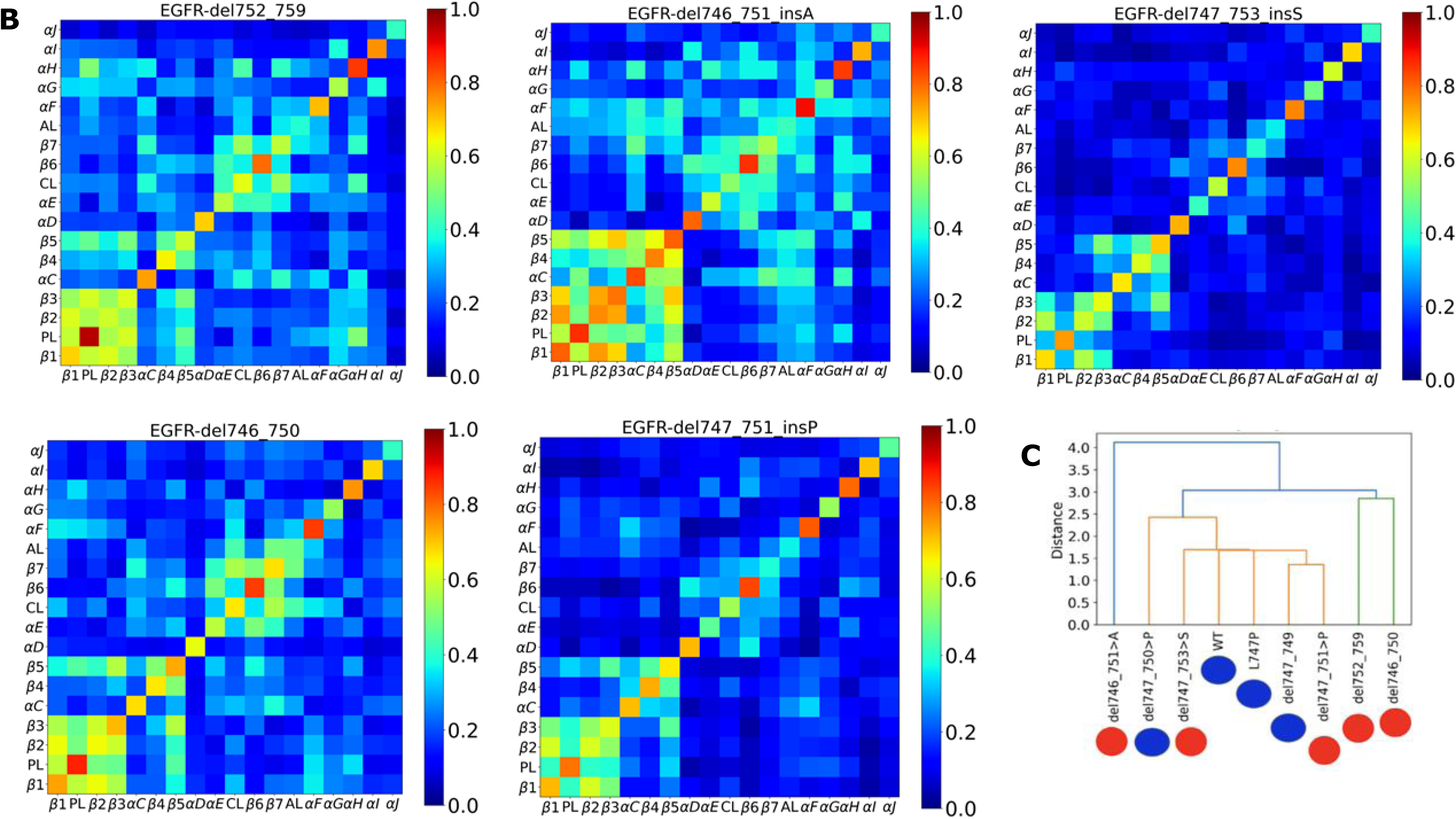
A. Coarse-grained dynamical cross-correlation matrices of EGFR Exon-19 deletion mutants from profile 1. B. Coarse-grained dynamical cross-correlation matrices of EGFR Exon-19 deletion mutants from profile 2. C. Hierarchical clustering of the EGFR Exon-19 deletion mutants. Mutations from profile 1 are marked in blue and mutations from profile 2 are marked in red.

### Hydrogen bond and solvent accessibility observed in unbiased simulations explain HDX data and provide validation of the observed changes in the protein conformational dynamics

To validate the molecular origins of the observed variations in the dynamics across profile 1 and profile 2 variant, we sought to compare the MD results with the experimentally observed HDX rates. The arrangement of the hydrogen bond network is a major determinant of the observed percent exchanges in HDX, as hydrogen bonds between protein residues must be broken to enable exchange of the hydrogen with deuterium in the solvent. Solvent accessibility also impacts HDX, however, as residues deeply buried in the protein have low rates of exchange due to lack of access to contact solvent molecules. As a measure of hydrogen bond ability, which should correlate with HDX based on these arguments, we plot “1 minus the hydrogen bond occupancy” from MD simulations of the EGFR deletion mutants as described in Methods. Comparing these results in Fig. 3A with the experimentally reported HDX results (Fig. S1C) supports the premise that hydrogen bond occupancy in MD is lower where HDX is higher.

**Figure 3.**
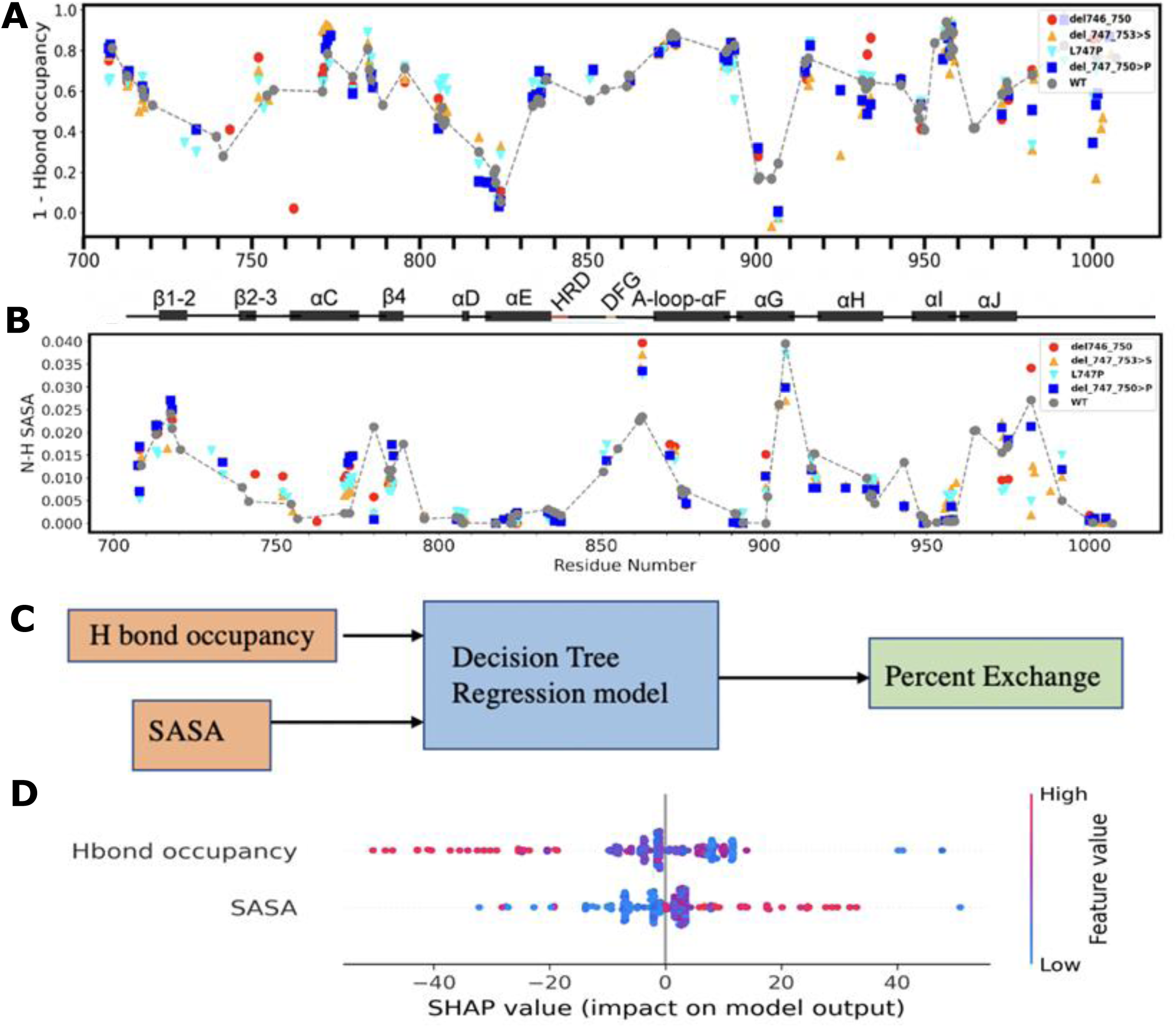
A. Hydrogen bond (H bond) occupancy per residue obtained from simulations. B. Solvent accessible surface area or SASA per residue obtained from simulations. C. Decision Tree regression model trained with H bond occupancy and SASA as inputs to predict Percent Exchange. D. SHAP values determine the contribution from the H bond and SASA towards % exchange, while also quantifying the nature of contribution. The length of the spread represents the predictive value and the direction represents the correlation. The results show that H bond occupancy holds a greater predictive value compared to the SASA. They also show that increased H bond occupancy will lead to lower exchange while increases SASA will lead to higher exchange.

To obtain a quantitative mapping between the computationally-determined H bond occupancy (Fig. 3A), solvent-accessible surface area (SASA) per residue (Fig. 3B), and the experimentally obtained percent HDX (Fig. S1C), we implement an information theory-based metric and a machine learning model. We use these to combine information in H bond occupancy and SASA to measure the explainability of the observed percent exchange in HDX studies. Across all systems analyzed, we find that combining SASA and H bond occupancy leads to an increase in the mutual information, i.e., more information on exchange is gained (see Table 3).

**Table 3:** Mutual information values of EGFR deletion mutants computed from molecular dynamics simulations.

| Protein | Mutual Information | Joint Mutual Information | Normalized Mutual Information |
| --- | --- | --- | --- |
| WT | 1.4 | 8.14 | 0.42 |
| L747P | 1.11 | 8.08 | 0.34 |
| $\Delta 747-750>P$ | 1.20 | 8.13 | 0.36 |
| $\Delta 747-753>S$ | 1.34 | 7.5 | 0.4 |
| $\Delta 746-750$ | 1.60 | 7.81 | 0.49 |

While it is challenging to find an equation that can combine H bond occupancy and SASA to provide % HDX as an output, machine learning models are ideal tools to combine multimodal data and can be trained to provide such a mapping. We trained a decision tree to take as input the H bond occupancy and SASA information as features and provide as output the % HDX (Fig 3C). Based on this decision tree regression model, we estimate the SHAP (SHapley Additive exPlanations) values (18), which measure the importance of each feature in the model in predicting its output while quantifying the nature of the contribution. Features with positive SHAP values positively impact the prediction, whereas those with negative values have a negative impact – with the magnitude of the SHAP value corresponding to the strength of the effect. As shown in Fig. 3D, H bond occupancy is more important towards predicting the % HDX than SASA, with higher H bond occupancy values (red) being associated with negative SHAP values; i.e. reducing the % HDX. Higher values for SASA (red), by contrast, are associated with positive SHAP values; i.e. with increased % HDX – consistent with our expectations and the results of Fig. 3A,B. Our results thus show that the observed dynamics captured in terms of principal motions are reflected in the observed exchange profiles in HDX experiments, thereby validating the computational findings.

### Conformational fluctuations coupled to solvation effects at long timescale explain the distinction between some of the EGFR Exon-19 deletion variants

Although we have investigated the molecular origins behind the observed differences in HDX for the different EGFR Exon 19-deleted variants based on the trajectories of limited timescale simulations, we note they are still not sufficient to fully explain the misclassified variants in the hierarchical cluster shown in Fig. 2C. In Fig. 4A, we observe that when % HDX is plotted against H bond occupancy for the wild type EGFR TKD, certain sequences fall in the lower left quadrant where both the H bond occupancy and % HDX simultaneously have low values (See Appendix). While this trend is counterintuitive, we note these same peptides also have low SASA (marked in red), which explains their unexpectedly low % HDX.

**Figure 4.**
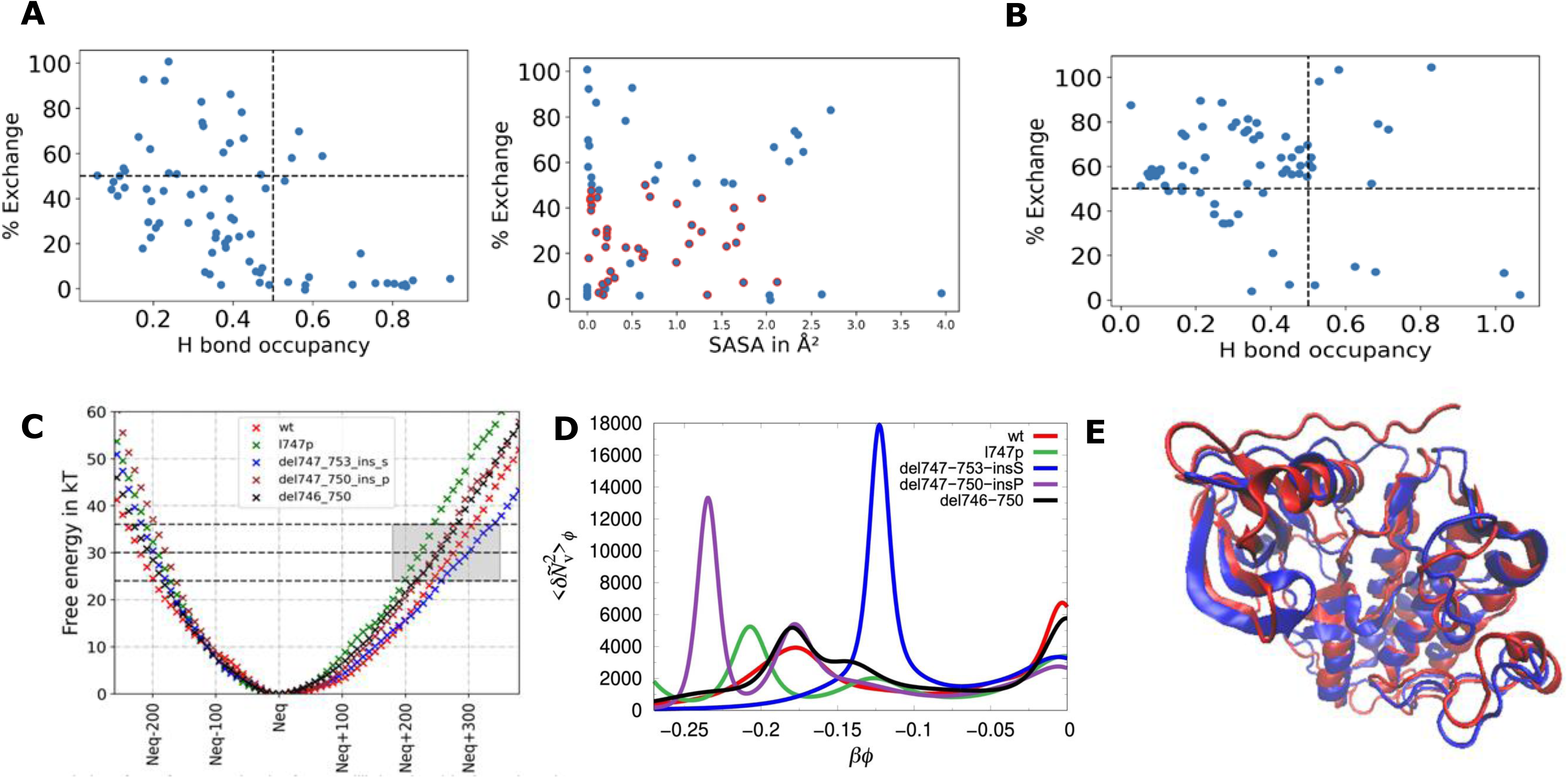
A. % exchange versus H bond occupancy and versus SASA for the peptides measured in HDX for wildtype EGFR. B. % exchange versus H bond occupancy and versus SASA for the peptides measured in HDX for del747-753-insS EGFR. C. Free energy profiles obtained through INDUS sampling. D. The water susceptibility function computed based on the fluctuations in the number of water molecules in the probe volume as a function of the applied field in the INDUS simulations (non-dimensionalized by **β**=1/k_B_T). E. Structural alignment between average structure for del-747-753-insS obtained from unbiased MD (shown in red) and INDUS corresponding to the activated state where the susceptibility function associated with water density function peaks (shown in blue). Structural alignment between average structure for del-747-753-insS obtained from unbiased MD (shown in red) and INDUS corresponding to an activated state conformation (shown in blue).

In the case of the Del747-753InsS system, one of the outliers in Fig. 2C, we find that certain sequences fall in under the upper right quadrant when % HDX is plotted against H bond occupancy (see Fig. 4B and Appendix), indicating high H bond occupancy and higher % HDX. This is trend is inconsistent with our intuition and is also not explained by the SASA values. We hypothesize that this is the result of limited timescale dynamics information provided by unbiased molecular dynamics, wherein the conformational sampling does not adequately represent partial unfolding of the backbone that could occur at timescales longer than the few microseconds captured by MD, but still within the experimental observation time of HDX experiments.

With this in mind, we computed the free energy change associated with solvent density fluctuations using Indirect Umbrella Sampling (INDUS) as depicted in Fig. 4C. We refer to conformations sampled under unbiased MD simulations as the equilibrium ensemble, corresponding to fluctuations near the minimum of the free energy landscape. In contrast, conformations sampled under INDUS bias at elevated free energy values (corresponds to the peak in the computed susceptibility function associated with collective solvent fluctuations, see below) are designated as activated states, representing rare but thermodynamically accessible conformational excursions coupled to pronounced solvent density fluctuations. In HDX studies, the protein is constantly undergoing transitions between partially folded and unfolded states, which is generally coupled with fluctuations in the density of water molecules in the probe volume around the protein. This is exactly the thermodynamic landscape explored in INDUS as depicted in Fig. 4C, D and E, where we compare the structures of the equilibrium conformation with that corresponding to the conformations where we see the peak in the susceptibility function associated with water density fluctuations (around free energy of 30 k_B_T)

We found that this mis-classified profile 2 system (Del747-753InsS) was more elastic with respect to movement of water in and out of the system compared with the other deletion variants, since the free energies associated with water density fluctuations were the lowest of all the EGFR deletion variants (Fig. 4D). As expected, a rise in the number of hydration waters, 〈*Ñ_v_*〉*_ϕ_* as we reduce the strength *ϕ* of the external biasing potential is observed for all the proteins studied. However, a sudden rise in 〈*Ñ_v_*〉*_ϕ_* at a certain *ϕ* for all the proteins signifies a collective transition. The solvation waters collectively respond to the external perturbation and increases suddenly which correlates with the conformational changes in the protein. Early appearance of a pronounced sudden jump in 〈*Ñ_v_*〉*_ϕ_* and the largest peak in the solvent susceptibility in Δ747-753InsS suggest a highest degree of solvent density fluctuation in this variant (Fig. 4E). Interestingly, this happens when the number of water molecules deviate from the equilibrium value in the rage of 200-400. Indeed, this onset in the peak in susceptibility corresponds to the activation free energy of ∼30 *k*_B_*T* indicating clearly that the timescale associated with such a collective solvent fluctuation is of the order of seconds as noted above.

We structurally aligned the average structure from INDUS (from the zone of 30 k_B_T) with respect to the equilibrium structure, and find the RMSD to be 4.51 A, suggesting only local changes in the secondary structure, and that the protein is not completely unfolding. To relate the free energy barrier to a characteristic transition timescale, we adopt an activated rate approximation based on Kramers theory (19). In this framework, the characteristic transition time can be estimated as 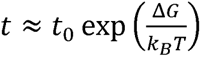, where *t*_0_ represents a molecular attempt time. For barriers in the range of ∼30 k_B_T, this expression yields transition timescales compatible with seconds, consistent with the experimental timescale probed in HDX-MS measurements. We can further explain the outlier results in Fig. 4B, wherein certain peptides that showed higher H bond occupancies also showed high % HDX. By extracting structures from the activated state of ∼30 k_B_T (gray box marked in Fig. 4C), we can compare the H bond profiles for the equilibrium structures with those corresponding to the activated state (Fig. 4E). The gray region in Fig. 4C corresponds to the activated free energy regime where solvent susceptibility peaks emerge, directly linking collective solvent density fluctuations to changes in hydrogen bond occupancy and local conformational rearrangements. We note that in the activated state, the Del747-753InsS system undergoes a loss in H bond occupancies in the outlier sequences. Fig. S2 depicts the difference in the H bond occupancy profile between the equilibrium and activated states; the outlier sequences for Fig. 4B are marked with a black bar, which show reduced H bond occupancies in the activated state but high occupancy in the equilibrium state. We note that the peak in susceptibility (Fig. 4D) for other systems occur at larger deviations of the number of water molecules from the equilibrium average and are associated with much higher free energy barrier indicating that those collective fluctuations are much slower than timescales accessed in HDX and hence are not physically realized in the experiments. Taken together, these results explain the paradoxical result of high H bond occupancy coupled with high exchange rates in Fig. 4B, namely that collective solvent fluctuations (as indicated by the peak in the susceptibility function in Fig. 4D) drive conformational fluctuations and alter H bond occupancies (Fig. 4E and S2) to influence exchange in HDX experiments. Moreover, they underscore the importance of long-timescale fluctuations in backbone conformations coupled to solvent density fluctuations which are important to consider in interpreting HDX studies.

## Discussion and Conclusion

In this work, we elucidate the impact of EGFR Exon-19 deletion mutations on kinase conformational dynamics by integrating MD simulations, machine learning, enhanced sampling and HDX-MS. We analyze MD simulations of protein dynamics using Principal Component Analysis and record distinct modes of conformational excursions for the EGFR deletion mutants in two TKI resistance ‘profiles’ identified experimentally. We observe that the motions in profile 1 (resistant) variants are largely localized around the nucleotide binding P-loop, the αC helix and the Activation loop, whereas for profile 2 (sensitive) variants the motions also extend to the N–lobe and C-lobe regions. We established a quantitative metric to support this qualitative observation by defining a dynamical cross correlation metric based on a coarse-grained representation of the residues into 18 sub-domains.

When binding to a prototypical kinase such as EGFR, ATP is bound at the interface between the C-lobe and the N-lobe (20). The ATP phosphates interact closely with the P-loop that connects strands β1 and β2 and also interact with a conserved lysine in strand β3. The N- and C-lobes are connected by a linker sequence referred to as the hinge that interacts with the adenine base of the ATP molecule through one or two H bonds (20, 21). Since profile 1 and profile 2 variants differ in their ATP binding affinities as estimated in *K*_M,ATP_ measurements, we wondered if the differences in conformational dynamics between the two profiles (Fig. 1) translates into changes in ATP binding. We therefore performed molecular docking studies across all of the clusters observed in the dynamics simulations. Our results indicate that indeed the ATP binding affinity relative to the binding affinity of erlotinib is a clear differentiator of profile 1 and profile 2 mutants, with profile 1 variants (L747P, Del747-750InsP) showing an increased relative affinity for ATP (Fig. 5). More generally, localized motions that do not induce relative conformational fluctuations between the C-lobe and the N-lobe preserve the order of the ATP binding site as seen in profile 1 variants. Largescale delocalized motion with relative conformational fluctuations between the two lobes as seen in profile 2 variants, however, induces disorder in the ATP binding site and negatively impacts ATP binding. This effect is felt in the P-loop region as well as the adjoining β1-β3 strands as clearly evidenced by the dynamical cross correlations between these sub-domains and those in the C-lobe (Fig. 2).

**Figure 5.**
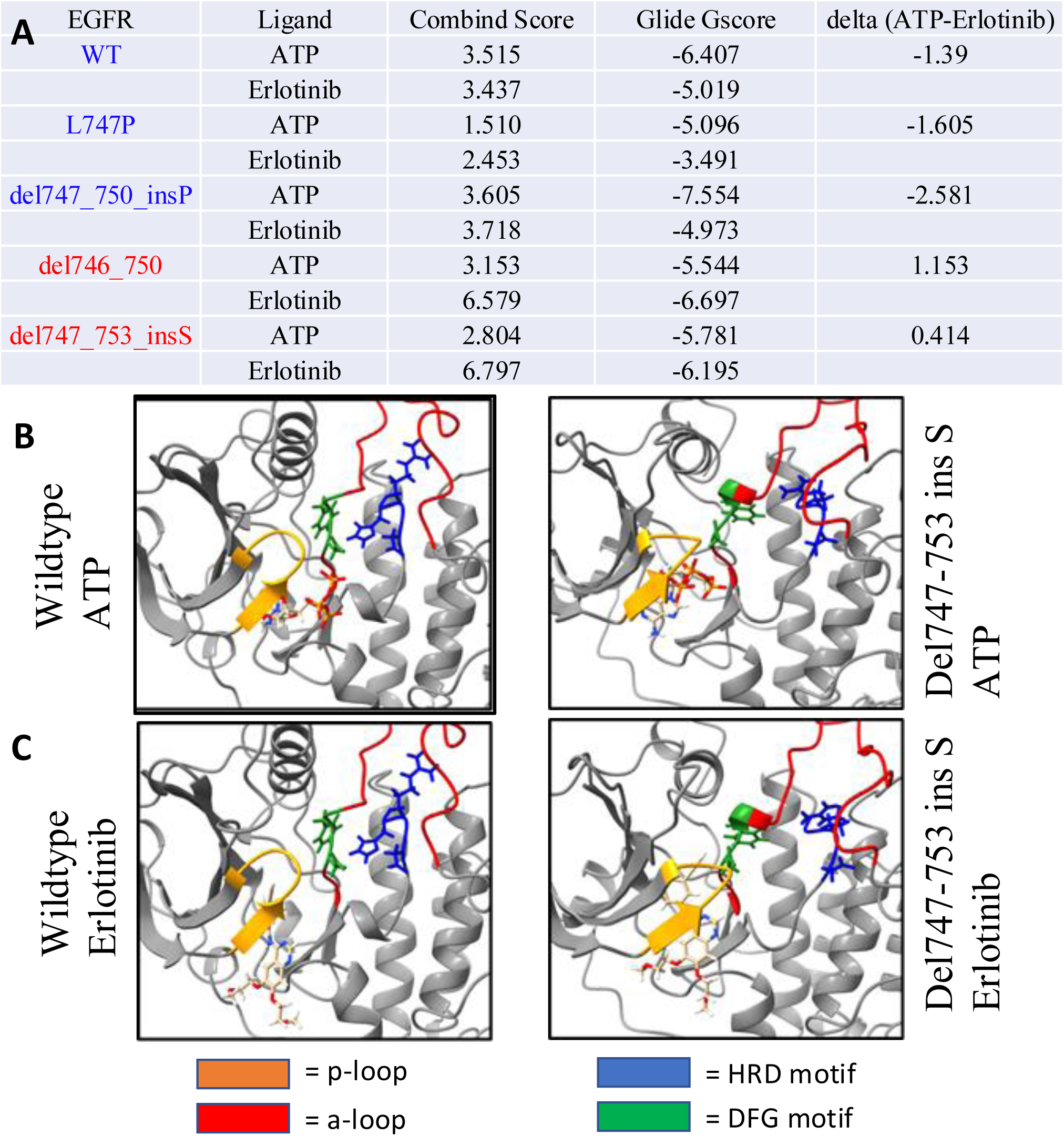
A. Results from molecular docking of ATP and erlotinib for EGFR variants. The zone that yielded the best docking score, which also conserves the known binding mode of the ligands is reported. B, C. Snapshots of docked ATP and erlotinib for representative profile 1 and profile 2 mutants are depicted.

By performing a pair-wise comparison of the subdomain-level dynamical cross correlations, we successfully classified the EGFR mutated systems through a hierarchical clustering analysis into profile 1 and profile 2 sets. Remarkably, all except two mutations, Del747-753InsS and Del747-751InsP were classified correctly with this approach. For the two misclassified mutants, we hypothesized a role for collective solvent fluctuations in shaping the overall protein dynamics in addition to the intrinsic-protein dynamics in a solvated environment, and we probed this further by analyzing our results in light of HDX experiments. Our studies also provided a machine learning-based interpretability metric, SHAP values, that quantify the contributions of hydrogen bonding and solvent accessibility in explaining the experimentally observed percentage exchanges in HDX. We conclude that, although H bond occupancy is the higher ranking metric in predicting HDX, combining it with SASA enhances prediction of the % HDX values. These observations led to the hypothesis that solvent accessibility coupled to long-timescale motion in protein conformational dynamics can play a role in exchange kinetics and could be important for our misclassification of the Del747-753InsS and Del747-751InsP variants. Hydration dynamics is an important component of protein dynamics but generally is a diverse timescale phenomenon and is therefore not typically well quantified when correlating MD studies with HDX. We implemented the INDUS enhanced sampling method to quantify the thermodynamic landscape associated with fluctuations in solvent density and find that the Del747-753InsS system has the lowest free energy barrier for invasion of solvent molecules into the probe volume, which is coupled to protein conformational flexibility. This result was further reinforced by examining the susceptibility which showed a peak in the excited state, thereby directly linking collective solvent density fluctuations to conformational fluctuations. Our results comparing the H bond and solvent accessibility in the ground and excited states directly demonstrate that conformation fluctuations coupled with solvent dynamics can contribute to exchange, and the exchange profiles match that of the HDX experiments.

Previous efforts integrating HDX-MS with molecular dynamics simulations have largely focused on interpreting exchange patterns in terms of equilibrium structural fluctuations and local flexibility (13, 14). In contrast, our approach extends this framework by incorporating enhanced sampling to capture rare solvent-coupled conformational excursions that bridge the timescale gap between microsecond MD trajectories and second-scale HDX exchange kinetics. By explicitly quantifying solvent susceptibility and its coupling to structural rearrangements, we provide a thermodynamic basis for understanding mutation-specific differences in exchange behavior.

We believe that integrating MD with INDUS and appending it with data science tools can be a powerful tool to improve the interpretability of HDX results and enable better mapping of molecular origins behind the observed deuterium exchanges of biomolecules studied in HDX. More importantly, such physics-based analysis involving enhanced sampling can be critical in explaining the origin of drug sensitivity and patient survival in cancer patients who present a varied genomic landscape by way of missense mutations and in-frame deletions and insertions.

## Methods

### Molecular Dynamics (MD) Simulations

We constructed variants of mutations in the EGFR structure (1M14 in protein data bank) using MODELLER (22). Homology modeling using MODELLER is a well-established approach for introducing sequence variants and reconstructing loop regions in kinase domains when mutant crystal structures are unavailable. In the present case, the deleted residues lie within flexible loop segments rather than conserved catalytic motifs, and local loop refinement was performed prior to molecular dynamics equilibration. The detailed protocol for loop refinement and assessment of the structures through the computation of several metrics including the discrete optimization of protein energy (DOPE) scores is provided in our earlier work (23). We note that for the structures modeled here, a comparison of the modeller and the AlphaFold2 generated conformations are practically the same as the anchor kinase conformations in the inactive state of the kinase are well represented in the protein data bank PDB. However, for active conformations, modeller provides the flexibility of the templates while providing a means to assess the quality of the structure, as we reported in a recent study (24).

Specifically, in addition to the wildtype, we studied L747P, DelL747-E749, DelL747-A750InsP, DelE746-A750, DelS752-I759, DelE746-T751InsA, DelL747-T751InsP, DelL747-P753InsS. To investigate the molecular underpinnings of observed differences in exchange profiles, unbiased molecular dynamics (MD) simulations totaling 4 microseconds (4 μs) were conducted using the Desmond simulation package (25) within the Schrödinger suite 2020-3, operating on a CentOS 7 Linux platform. Simulations were performed in the NPT ensemble at 300 K and 1 bar pressure, employing the Optimized Potentials for Liquid Simulations (OPLS 2005) force field (26, 27). The protein complex was centered in an orthorhombic box, and TIP3P water molecules, along with buffer atoms, were filled within a 10 Å distance from protein atoms to the box’s edge. To neutralize the system, counter ions (Na+ and Cl−) were added randomly, resulting in a salt concentration of 150 mM. Long-range electrostatic interactions were computed using the particle mesh Ewald method (28, 29), with a Coulomb interactions cutoff radius of 9 Å. Non-bonded forces were integrated using the r-RESPA integrator (30). Temperature and pressure control were achieved using the Nose Hoover chain coupling with a 1.0 ps relaxation time at 300 K, and the Martyna-Tobias-Klein chain isotropic coupling barostat method (31) with a 2.0 ps coupling constant for pressure control. The protocols employed above have been previously employed in studying EGFR TKD mutations (32, 33).

### Data Analysis

Trajectory analyses were performed, unless otherwise noted, on the whole trajectory using the MDAnalysis package (34) in Python. Python scripts that implement these analyses are available on GitHub (35). Trajectories were saved at intervals of 100/200 ps for subsequent analysis, including examination of polar interactions and root mean square deviations (RMSD) to monitor simulation stability. Principal Component Analysis (PCA) was performed on the trajectories using the GROMACS, projecting the results onto the top two components after removing center of mass translation and rotation. Python scripts that implement these analyses are available on GitHub. Trajectories were saved at intervals of 100/200 ps for subsequent analysis, including examination of polar interactions and root mean square deviations (RMSD) to monitor simulation stability. Covariance matrix of atomic coordinates are constructed and then diagonalized to find the top two principal components (termed PC1 and PC2) corresponding to the top two largest eigen values. Hydrogen bond computation and solvent accessible surface area calculations were conducted using GROMACS.

#### Dynamical Cross Correlation (DCC)

The normalized covariance matrix gives the traditional residue-level dynamical cross correlation (DCC). given by:

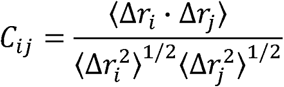

here, Δ*r_i,k_* = *r_i,k_* − *μ_r,i,_*, where *r_i,k_* is coordinate of residue *i* in frame *k*; and *µ_r,i_* is the mean location of residue *i* over total *N* frames. <u>Coarse-grained DCCs</u>: to obtain correlations at the sub-domains or secondary structure level and ascribe a broad mechanism, we coarse-grained the residue-level DCC matrix to their secondary structure (36) as follows. The following residue number ranges were assigned to each of the 18 subdomains (see Table 1). The NxN BW-DC matrix was reduced to a 18×18 coarse-grained matrix by averaging the correlation coefficients in each sub-block.

The H bond occupancy per residue was computed based on the number of frames a residue has a hydrogen bond / total number of frames. The H bond occupancy of the peptide was obtained using the mean of H bond occupancy over the residues of the peptides. The SASA per residue was computed for N and H of each residue and the SASA per peptide was taken as the mean of this SASA per residue for the residues in a peptide.

From information theory, the mutual information between two variables is defined as I(Z; X) = H(Z) - H(Z|X) (37). I(Z) represents the reduction in the uncertainty of Z due to information on X; in our implementation, Z = % exchange in HDX and X is the H bond occupancy. H(Z) is defined as H(Z) = - P(Z)log(P(Z)), where P(Z) is the distribution of Z. Similarly, joint mutual information is defined I(X, Y; Z), where Z= % exchange and X= H bond occupancy and Y is the SASA and I(X, Y; Z) is defined as the information that is gained in Z by knowing X and Y. We compute mutual information, joint mutual information and the normalized mutual information and tabulate them for EGFR deletion systems (Table 3).

### Indirect Umbrella Sampling (INDUS) Simulations

We compute the free energy change associated with solvent density fluctuations in the solvation shell of protein using the Indirect Umbrella Sampling (INDUS) (15)technique. We have performed all our umbrella sampling simulations using GROMACS 4.5.3 patched with INDUS. The hydration of the protein was modulated in INDUS by considering a probe volume *v*, corresponding to its hydration shell. The probe volume *v* is defined as the union of spherical sub-volumes which are pegged into every protein surface heavy atom. Each sub-volume has a radius of *R_v_* = 0.6 nm to capture the first layer of hydration waters. Note that the size and shape of the probe volume dynamically changes in in response to the conformational fluctuations of the protein. The obtained free energy from INDUS simulations was unbiased using the weighted histogram analysis method. Trajectories were processed in Python for further analysis, and EGFR structure mutations were introduced using MODELLER. All scripts and codes utilized in these analyses are publicly available on GitHub (35).

We employed the indirect umbrella sampling (INDUS) technique to investigate the free energetics of water intrusion and displacement from the solvation shell of the wild-type protein and its 4 different mutations. The INDUS method allows modulating the number of water molecules, *N_v_*, within the solvation shell of volume *v* of a protein by introducing a bias on a coarse-grained water number *Ñ_v_*. Unlike *N_v_*, *Ñ_v_* represents a continuous variable of particle positions and does not induce abrupt forces upon application of biasing potential (16, 17, 38–41). The observation volume *v* for the INDUS simulation is defined as the combination of sub-volumes with radius *r_v_* = 0.6 nm, each bound to all the heavy atoms of proteins. The collective assembly of these sub-volumes approximately depicts the solvation shell of the protein. A simulation snapshot of the WT protein, highlighting the observation volume *v* is shown in Figure S3 A. It should be noted that the observation volume roughly mimics the shape of the protein and can dynamically adjust with changes in its conformation (16, 41).

A biasing potential of the form 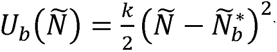 with force constant *k* is employed for each protein, as described in Table 4. We first performed an unbiased simulation for 10 ns to compute the variance in the number of water molecules (〈*δN*^2^〉_0_) within the observation volume of a specific protein (16, 17, 38–41). The value of k is selected such that it is at least four times greater than 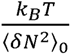, where *k*_B_ is the Boltzmann constant, and *T* is the temperature at which the simulation is performed. The final configuration of the unbiased simulation is utilized for biased simulation, with the biasing number of water molecules ranging from 1700 to 2600 with intervals of 50 (Table 4). The duration of simulation for each biasing window is taken to be 10 ns. We used the weighted histogram analysis method (WHAM) to combine the biased probability distributions to estimate the free energy *βF_v_*(*Ñ*) = −ln (*P_v_*(*Ñ*), where *P_v_*(*Ñ*) is the probability of observing *Ñ* water molecules in *v* and *β* = 1/*k_B_T*. Estimated *F_v_*(*Ñ*) is then reweighted to investigate how a linear biasing potential such as *U_ϕ_*≡ *ϕÑ_v_*, modulate the solvation of the proteins (39). *ϕ* parameter is the strength of the biasing potential applied to *Ñ_v_* coarse-grained water in volume *v*. The average number of waters 〈*Ñ_v_*〉*_ϕ_* as a function of *βϕ* is then calculated using the biased Hamiltonians, ℌ*_ϕ_ =* ℌ_0_ + *U_ϕ_*, where ℌ*_ϕ_* is the Hamiltonian in absence of *ϕ*. We first estimate the biased probability 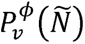 using the unbiased probability *P_v_*(*Ñ*) using 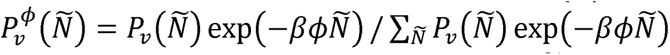. This biased probability is then used to estimate average number of waters such as 〈*Ñ_v_*〉*_ϕ_* = ∑*_Ñ_ ÑP_v_*(*Ñ*) and its susceptibility 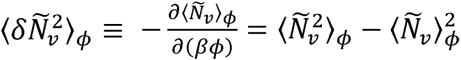. As mentioned earlier, the linear potential *U_ϕ_* ≡ *ϕÑ_v_* influence the average number of waters 〈*Ñ_v_*〉*_ϕ_* in the probe volume. When *ϕ*(or *βϕ*) = 0 there is no linear biased applied and the equilibrium number water within the probe volume is not modulated. On the other hand, upon application of negative biasing potential (*ϕ*(or *βϕ*) < 0) a greater number of waters are pushed to the solvation shell as compared to the number of waters present in the equilibrium states. Here, *β* = 1/*k_B_T* and *ϕ* controls the strength of the external perturbation applied to the coarse-grained water number within the probe volume. Physically, *βϕ* therefore represents the magnitude of the thermodynamic bias used to modulate hydration fluctuations, enabling us to quantify the system’s susceptibility to collective solvent density changes. *ϕ* parameter decides the strength of the biasing potential applied to *Ñ_v_* coarse-grained water in volume *v*. The average number of waters 〈*Ñ_v_*〉*ϕ* and its susceptibility 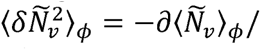/∂(*βϕ*) as a function of *βϕ* is collected and plotted (see Fig. S3 B and Fig. 4D).

**Table 4:** Parameters related to the INDUS enhanced sampling simulations for EGFR variant systems.

| Protein | Spring constant, k<br>(kJ Mol<sup>-1</sup>) | $N_b^*$ for biasing<br>windows | Biasing<br>simulation length<br>(ns) per window | Average number of<br>waters in v under<br>no bias ($\langle \tilde{N}_v \rangle_{\phi=0}$) |
| --- | --- | --- | --- | --- |
| WT | 0.012 | 1700 to 2600 with<br>$dN_b^* = 50$ | 10 | 2163 |
| L747P | 0.015 | 1700 to 2600 with<br>$dN_b^* = 50$ | 10 | 2074 |
| $\Delta 747-750 > P$ | 0.010 | 1700 to 2600 with<br>$dN_b^* = 50$ | 10 | 1967 |
| $\Delta 747-753 > S$ | 0.0111 | 1700 to 2500 with<br>$dN_b^* = 50$ | 10 | 2030 |
| $\Delta 746-750$ | 0.015 | 1700 to 2600 with<br>$dN_b^* = 50$ | 10 | 2103 |

Since the different EGFR Exon-19 deletion mutant have a different numbers of water molecules in the probe volume at equilibrium (N_eq_), we shift the plots so that the x-axis represents the deviation of the water molecules in the probe volume from the equilibrium value, N_eq_. The y-axis is also shifted so that the minimum corresponds to zero free energy. Because each EGFR variant exhibits a distinct equilibrium hydration level within the probe volume, subtracting N_eq_ ensures that N − N_eq_ = 0 corresponds to the unbiased equilibrium condition for each system. This normalization allows direct comparison of hydration fluctuations across variants without conflating intrinsic differences in baseline hydration with solvent response to bias.

### Molecular Docking

#### Preparation of protein and ligand structures

The ligands were prepared by adding missing atoms, assigning charges, and optimizing the geometry following standard protocol in Schrodinger’s LigPrep package. The EGFR receptor structures were prepared by adding missing atoms, assigning charges, optimizing the hydrogen bonding network, and performing restrained minimization following standard protocol using Schrodinger’s Protein Preparation Wizard tool. The receptor structures were then aligned to a known crystal structure of the EGFR (PDB: 1M17) tyrosine kinase domain in complex with erlotinib. The centroid of erlotinib is used as the center for the docking grid, as this the best estimate of EGFR’s binding site.

#### Simulation

Schrodinger’s (v. 2019.4) Glide software was used to performing docking simulations for each of the cluster identified through PCA (42, 43). Ligands were docked using default Glide SP settings, except that “Enhanced Sampling” was set to 4, to increase the number of ligand conformers considered. For each ligand, a up to 100 of the most highly ranked poses were considered for further analysis.

#### Prediction of Binding Pose and Affinity using ComBind

The poses outputted by the Glide software package (Fig. 5 B, C) are scored using the internal GlideScore function, traditionally used as an estimate of binding affinity. Studies have shown that this internal score is actually a poor estimate of binding affinity, which can lead to inaccurate prediction of binding pose. Recently, new methods have been created, combining information from both physics-based and ligand-based approaches to provide better estimates of binding affinity. One such method, ComBind (44), additionally considers other ligands known to bind the target protein (whose binding poses are not known), resulting in more accurate predictions. The ComBind methodology is used in our study for the prediction of the binding pose and affinity of our ligands to the EGFR systems. First, a set of diverse ligands known to bind EGFR were compiled.

The helper ligands (see Appendix) and candidate ligands (ligands of interest) are individually docked to the target EGFR systems using the simulation protocol previously described. Combind is used to predict poses for all of the helper ligands. We omit the “substructure similarity” term in calculation of Combind scores given the diverse range of high affinity binders chosen for our set of helper ligands. For each candidate ligand, a pose is selected that minimizes the Combind score with respect to the helper ligands. The Combind score of each candidate ligand in its predicted pose is used as a prediction of its binding affinity to the target EGFR system. Of note, a more positive Combind score equates to a higher binding affinity. The pose that yielded the lowest (most negative) docking score that also preserved the known modes of binding was recorded and reported. The detailed procedure for identifying know binding modes is described in earlier works (21, 45), which was closely followed here.

## Supplementary Figures

**Figure S1.** A. Exon-19 deletion mutations (deletion residues) fall in the loop between beta3 strand and αC helix. B. Note that in the figure we have used the shorthand notation Δ instead of Del to save space. The EGFR Exon-19 deletion mutations fall in two profiles based on ATP binding affinity (shown in blue and red). Michaelis-Menten plots for the indicated EGFR kinase domains in the presence of 20 μM peptide substrate (n = 3 biological replicates, each in technical triplicate). Normalized initial velocity data were plotted against ATP concentration (error bars represent SD across replicates) and fit to the Michaelis-Menten equation to obtain values for *K*_M, ATP_. The mean value of *K*_M, ATP_ across all replicates is plotted (± SD). Data for wild type kinase are shown in black, profile 1 variants in blue, profile 2 variants in red, L858R variant in olive, and the inactivated DelL747-E749 variant in gray. C. Percentage Exchange obtained from HDX for EGFR variants. Plot of the percent exchange of backbone amide hydrogens from HDX-MS experiments. DelL747-A750InsP (blue square), L747P (cyan triangle), DelE746-A750 (red circles), and DelL747-P753InsS (orange triangles) variants in the absence of ATP or inhibitor. Mean percent exchange (± SD) for at least 2 biological repeats (3 technical repeats/n) is plotted against the median residue number of the peptide in wild type EGFR numbering. Positions of secondary structure elements are denoted at the top of the figure, with functionally important regions (e.g., αD, HRD motif, and activation loop) indicated. Wild type data are represented by gray diamonds and two dashed gray lines representing the range (± SD) for each point.

**Figure S2.** Hydrogen bond network alteration at long timescales for the peptides that were outliers (i.e., belonging to upper right quadrant in Fig. 4B), marked in black. The top panel depicts the H bond occupancies per residue for the equilibrium structure. The middle panel depicts the H bond occupancy for the structure from the activated state in the INDUS simulations induced by collective solvent fluctuations (structures corresponding to the gray box in Fig. 4C). The bottom panel reports the difference in the two occupancy between middle and top panels to highlight the difference in H bond occupancy due to collective solvent fluctuations. The regions showing large differences in the H bond occupancy also overlap with those identified in HDX as regions with large exchange. The experimental data are derived from (9).

**Figure S3:** A. A simulation snapshot of the WT protein illustrating the observation or probe volume, *v* for INDUS simulations. The water molecules that are within this observation volume are shown licorice and the rest as lines. The protein atoms are shown in space-filled representation colored according to their residue types. B. Average number of water molecules in the probe volume as a function of the applied field in the INDUS simulations (non-dimensionalized by **β**=1/k_B_T). The experimental data are derived from (9).

## Supplementary Movies SI-M1 to SI-M9

**SI-M1:** Depicts movies of associated domain motions extracted from principal components (PCs) summarized in Table 1 for WT. Projection along PC1 is shown in A and that along PC2 in B.

**SI-M2:** Depicts movies of associated domain motions extracted from principal components summarized in Table 1 for L747P. Projection along PC1 is shown in A and that along PC2 in B.

**SI-M3:** Depicts movies of associated domain motions extracted from principal components summarized in Table 1 for del 747_750_insP. Projection along PC1 is shown in A and that along PC2 in B.

**SI-M4:** Depicts movies of associated domain motions extracted from principal components summarized in Table 1 for del 747_749. Projection along PC1 is shown in A and that along PC2 in B.

**SI-M5:** Depicts movies of associated domain motions extracted from principal components summarized in Table 1 for del 746_750. Projection along PC1 is shown in A and that along PC2 in B.

**SI-M6:** Depicts movies of associated domain motions extracted from principal components summarized in Table 1 for del 747_753_insS. Projection along PC1 is shown in A and that along PC2 in B.

**SI-M7:** Depicts movies of associated domain motions extracted from principal components summarized in Table 1 for del_746_751_insA. Projection along PC1 is shown in A and that along PC2 in B.

**SI-M8:** Depicts movies of associated domain motions extracted from principal components summarized in Table 1 for del 752_759. Projection along PC1 is shown in A and that along PC2 in B.

**SI-M9:** Depicts movies of associated domain motions extracted from principal components summarized in Table 1 for del 747_751_insP. Projection along PC1 is shown in A and that along PC2 in B.

## Appendix

The peptide sequences that fall under the lower left quadrant of the WT percent exchange vs. H bond occupancy (Fig 4A):

[’LRILKETEFKKIKVLGSGA’,

’RILKETEF’,

’VMASVDNPHVCRL’,

‘SVDNPHVCRL’,

’LGICL’,

’LTSTVQL’,

’TSTVQL’,

’VQLITQL’,

’MPFGCL’,

LDYVREHKDNIGSQYL’,

’YLEDRRLVHRDLAA’,

’LEDRRLVHRDLAARN’,

’LEDRRLVHRDLAARNVL’,

’RRLVHRDLAARNVL’,

’VLVKTPQHVKITDFGL’,

’VKTPQHVKITDFGLAKLLGAE’,

’KEYHAEGGKVPIKWMAL’,

’YHAEGGKVPIKWMAL’,

’ESILHRIYTHQSDVW’,

’ESILHRIYTHQSDVWS’,

’HRIYTHQSDVWS’,

’ISSILEKGERLPQPPICT’,

’SILEKGERLPQPPICT’,

’ILEKGERLPQPPIC’,

’ILEKGERLPQPPICT’,

’EKGERLPQPPICT’,

’IDVYM’,

’IMVKC’,

’IMVKCWM’,

’WMIDADSRPKFREL’,

’MIDADSRPKFRE’,

’MIDADSRPKFREL’,

’IDADSRPKFRE’,

’IDADSRPKFREL’,

’DADSRPKFRE’,

’FSKMARDPQRY’,

’MARDPQRY’,

’ARDPQRY’

The peptide sequences that fall under the upper right quadrant of the del747-753-insS percent exchange vs. H bond occupancy (Fig 4B):

[’DYVREHKDNIGSQ’,

’DYVREHKDNIGSQYLL’,

’IAKGMNY’,

’ISSIL’,

’ISSILEKGERLPQPPICT’,

’LVIQGDE’,

’YRALMDE’,

’YRALMDEEDM’,

’YRALMDEEDMD’

List of these 22 “helper” ligands: gefitinib, zorifertinib, sapitinib, poziotinib, tarlox, cln081, az5104, mobocertinib, lapatinib, icotinib, neratinib, dacomitinib, alomunertinib, olmutinib, pyrotinib, brigatinib, vandetanib, mavelertinibnazartinib, abivertinib, rociletinibnaquotinib, lazertinib, alflutinib.

## Notes

https://github.com/KesPatil/EGFR_kinase

### Competing Interest Statement

The authors have declared no competing interest.

### Summary of Updates

Methods, Results interpretation and clarification, Author list were updated. Summary - minor changes overall.

### Funding

- National Cancer Institute: CA250044

