## Supplementary figures and images for "Coupled Solvent and Protein Dynamics Confer Differences in Exon-19 Deletion Mutants of the Epidermal Growth Factor Receptor Kinase"

### KP-exon19-SI-figures.pdf

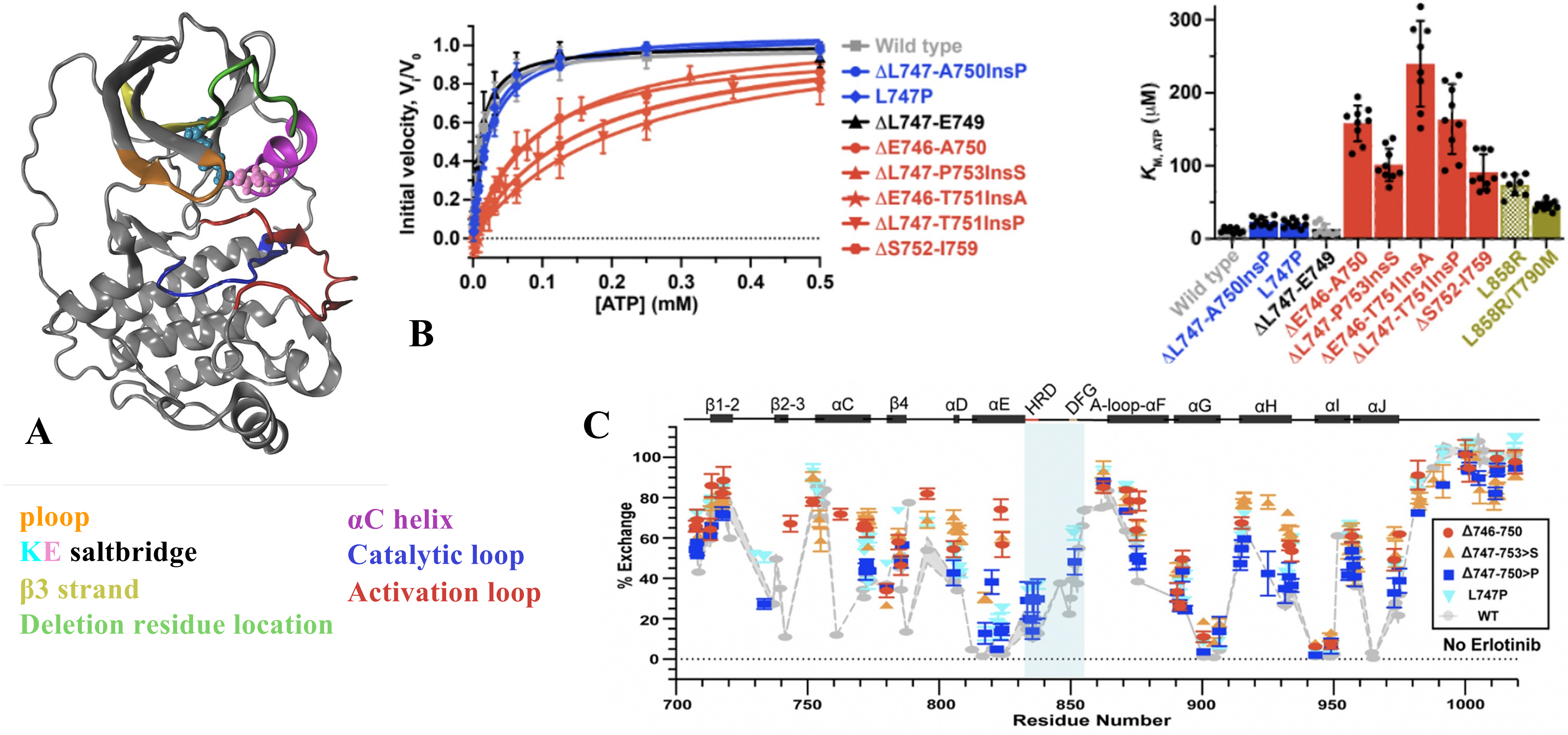

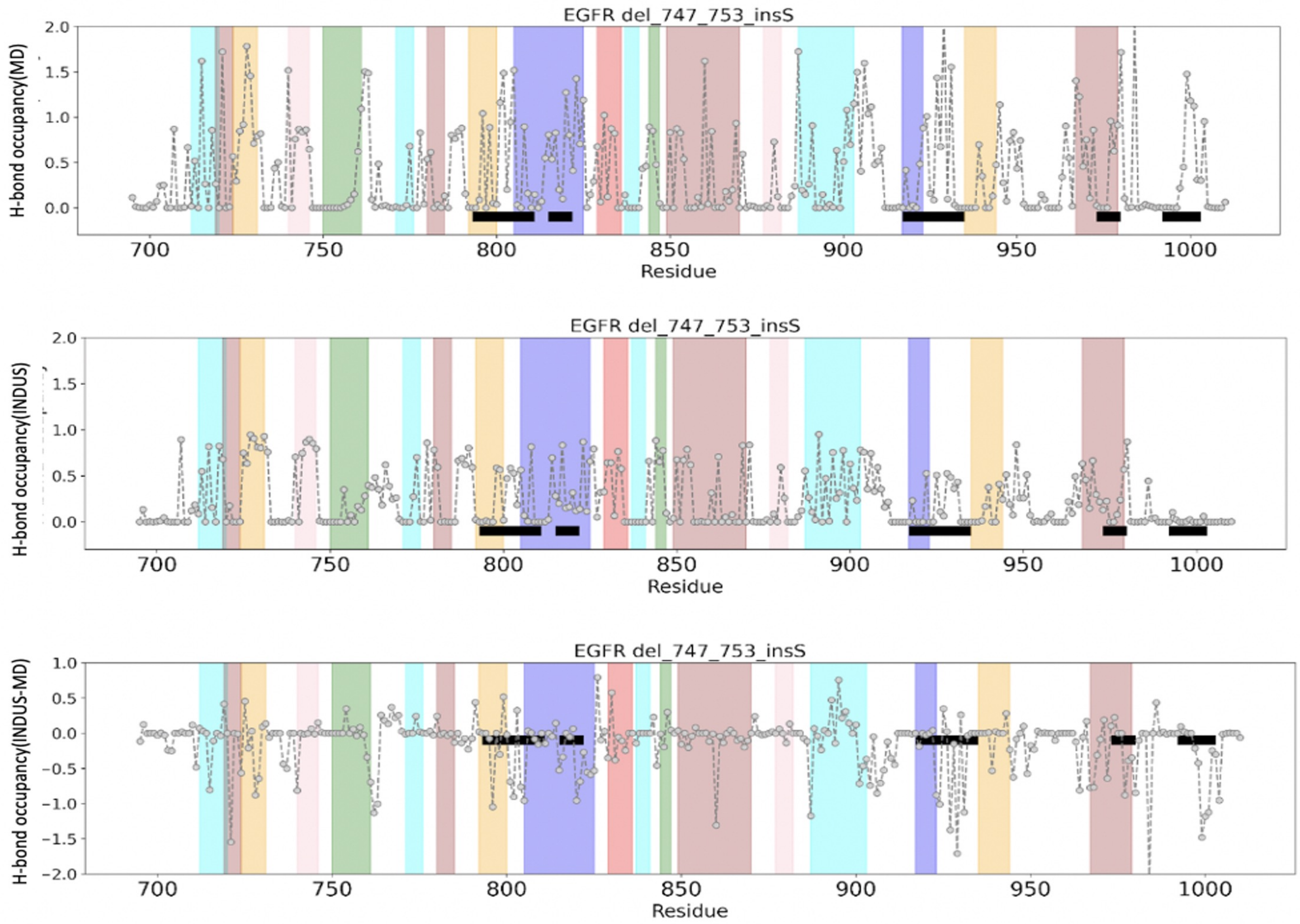

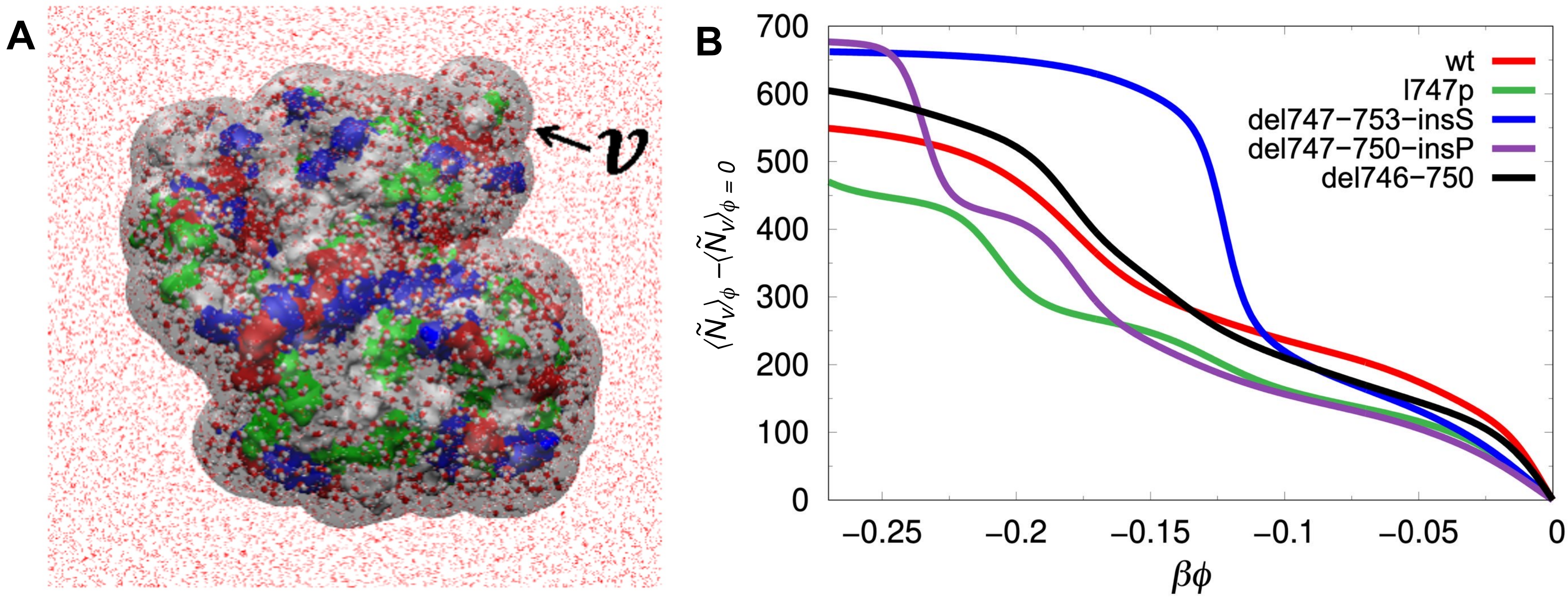
